# Antimicrobial prodrug activation by the staphylococcal glyoxalase GloB

**DOI:** 10.1101/2020.07.23.214460

**Authors:** Marwa O. Mikati, Justin J. Miller, Damon M. Osbourn, Naomi Ghebremichael, Ishaan T. Shah, Carey-Ann D. Burnham, Kenneth M. Heidel, Victoria C. Yan, Florian L. Muller, Cynthia S. Dowd, Rachel L. Edwards, Audrey R. Odom John

## Abstract

With the rising prevalence of multidrug-resistance, there is an urgent need to develop novel antibiotics. Many putative antibiotics demonstrate promising *in vitro* potency but fail *in vivo* due to poor drug-like qualities (e.g. serum half-life, oral absorption, solubility, toxicity). These drug-like properties can be modified through the addition of chemical protecting groups, creating “prodrugs” that are activated prior to target inhibition. Lipophilic prodrugging techniques, including the attachment of a pivaloyloxymethyl group, have garnered attention for their ability to increase cellular permeability by masking charged residues and the relative ease of the chemical prodrugging process. Unfortunately, pivaloyloxymethyl prodrugs are rapidly activated by human sera, rendering any membrane permeability qualities absent during clinical treatment. Identification of the bacterial prodrug activation pathway(s) will allow for the development of host-stable and microbe-targeted prodrug therapies. Here, we use two zoonotic staphylococcal species, *S. schleiferi* and *S. pseudintermedius*, to establish the mechanism of carboxy ester prodrug activation. Using a forward genetic screen, we identify a conserved locus in both species encoding the enzyme hydroxyacylglutathione hydrolase (GloB), whose loss-of-function confers resistance to carboxy ester prodrugs. We enzymatically characterize GloB and demonstrate that it is a functional glyoxalase II enzyme, which has the capacity to activate carboxy ester prodrugs. As GloB homologs are both widespread and diverse in sequence, our findings suggest that GloB may be a useful mechanism for developing species-or genus-level prodrug targeting strategies.

## INTRODUCTION

In 2019, the United States recorded 2.8 million antibiotic-resistant infections, resulting in over 35,000 deaths (1). The recent surge in antibiotic use in the setting of the COVID-19 pandemic portends an acceleration of the antibiotic resistance threat (1, 2). *Staphylococcus aureus* is a formidable human pathogen that causes a wide variety of invasive and life-threatening infections. Closely related staphylococcal species, *S. pseudintermedius* and *S. schleiferi*, cause similar skin, soft tissue, and invasive infections in companion animals and are increasingly appreciated as serious pathogens of humans (3–6). Rising rates of methicillin resistance are reported in all three species, with methicillin-resistant *S. aureus* (MRSA) labeled a “serious threat” by the Centers for Disease Control and Prevention (CDC) (7–10). Novel antimicrobial strategies that circumvent existing drug resistance mechanisms are urgently needed.

Bacterial metabolism is a promising area for antimicrobial development (11, 12). Many metabolic processes are essential for bacterial growth and pathogenesis. However, targeting metabolic processes can be inherently challenging, as a substantial portion of metabolism involves the catalytic transformation of highly charged substrates (e.g. phosphate transfer reactions). Substrate-competitive inhibitors of metabolic enzymes frequently deploy phosphonate functional groups as isosteric phosphate mimics (13). These negatively charged phosphonate antimetabolite inhibitors are prone to unacceptable drug-like characteristics and often diffuse poorly across membranes (14–19).

Prodrugging, or the modification of an inhibitor through addition of labile chemical adducts, is a common medicinal chemistry strategy to improve drug-like properties of an inhibitor under development (19–21). As promoieties are released prior to inhibitor-target engagement, prodrugging can temporarily cloak problematic pharmacokinetic properties such as poor absorption or solubility. For example, the third-generation cephalosporin, cefditoren, is poorly absorbed in the small intestine unless the phosphonate residues are masked with a lipophilic pivaloyloxymethyl (POM) promoiety, in the form of cefditoren pivoxil (22). Similarly, nucleoside analogues are generally cell-impermeable, but their cognate prodrugs have much improved cellular penetration and antiviral efficacy, as seen in remdesivir (SARS-CoV2), tenofovir disoproxil (HIV), and sofosbuvir (hepatitis C virus, HCV) (23–26). We have recently employed lipophilic prodrugging strategies to increase the efficacy of broad-spectrum antimicrobial phosphonate antibiotics. Notably, POM ester modification of a phosphonate isoprenoid biosynthesis inhibitor (ERJ) increases antistaphyloccal activity by 200- and 500-fold for *S. schleiferi* and *S. pseudintermedius*, respectively (27). Similar dramatic potency gains are observed for the same class of compounds against *Mycobacterium tuberculosis, Yersinia pestis, Franciscella novicida*, and the malaria parasite, *Plasmodium falciparum* (16, 28–31).

While pivaloyloxymethyl ester (POM)-prodrugs demonstrate remarkable potency *in vitro*, POM-promoieties are known to be rapidly hydrolyzed by serum carboxylesterases (32). If cell-impermeable phosphonate antibiotics are to be effective at the site of infection, the promoiety must be resistant to premature bioactivation during absorption and distribution in the circulation. This specificity in prodrug activation has been successfully achieved for liver-targeted prodrugs, using the “HepDirect” prodrug approach, but has not yet been deployed for antibiotic delivery. HepDirect prodrugs are cleaved via a hepatocyte-specific cytochrome P450 enzyme, CYP3A4, and are resistant to cleavage by other human esterases (33). Selective bioactivation of prodrugs within microbes would not only increase the circulating half-life, but may also improve the therapeutic selectivity of therapeutics that target microbial enzymes with human homologs. Understanding the molecular basis of host and microbe prodrug activation will facilitate design of microbially targeted prodrugs.

In this study, we use two zoonotic staphylococcal species, *S. schleiferi* and *S. pseudintermedius*, to uncover the enzymatic mechanism of prodrug activation in staphylococci. We identify and characterize the first bacterial carboxy ester prodrug activating enzyme, GloB, a type II glyoxalase. Using detailed biochemical analyses, we demonstrate that GloB recognizes the carboxy ester portion of the prodrug and is responsible for prodrug activation. Since GloB homologues are broadly maintained, yet have substantial sequence variation, we propose that this group of enzymes may be a strategy towards microbe-specific prodrug targeting.

## Results

### Selection of prodrug-resistant staphylococci

In our previous study, we identified phosphonate antibiotics with activity against zoonotic staphylococci (*S. schleiferi* and *S. pseudintermedius*) (27). Lipophilic carboxy ester prodrug modification of these phosphonates dramatically increases antistaphylococcal potency, presumably through increased cellular penetration (Fig. 1A, B). However, prodrug modifications block direct engagement of inhibitors with their enzyme target (27). For this reason, we hypothesized that one or more staphylococcal esterases were required for intracellular prodrug activation (Fig. 1A). To identify candidate prodrug activating enzymes, we designed a genetic screen/counter-screen strategy to enrich for staphylococcal strains that fail to activate lipophilic ester prodrugs.

**Figure 1.**
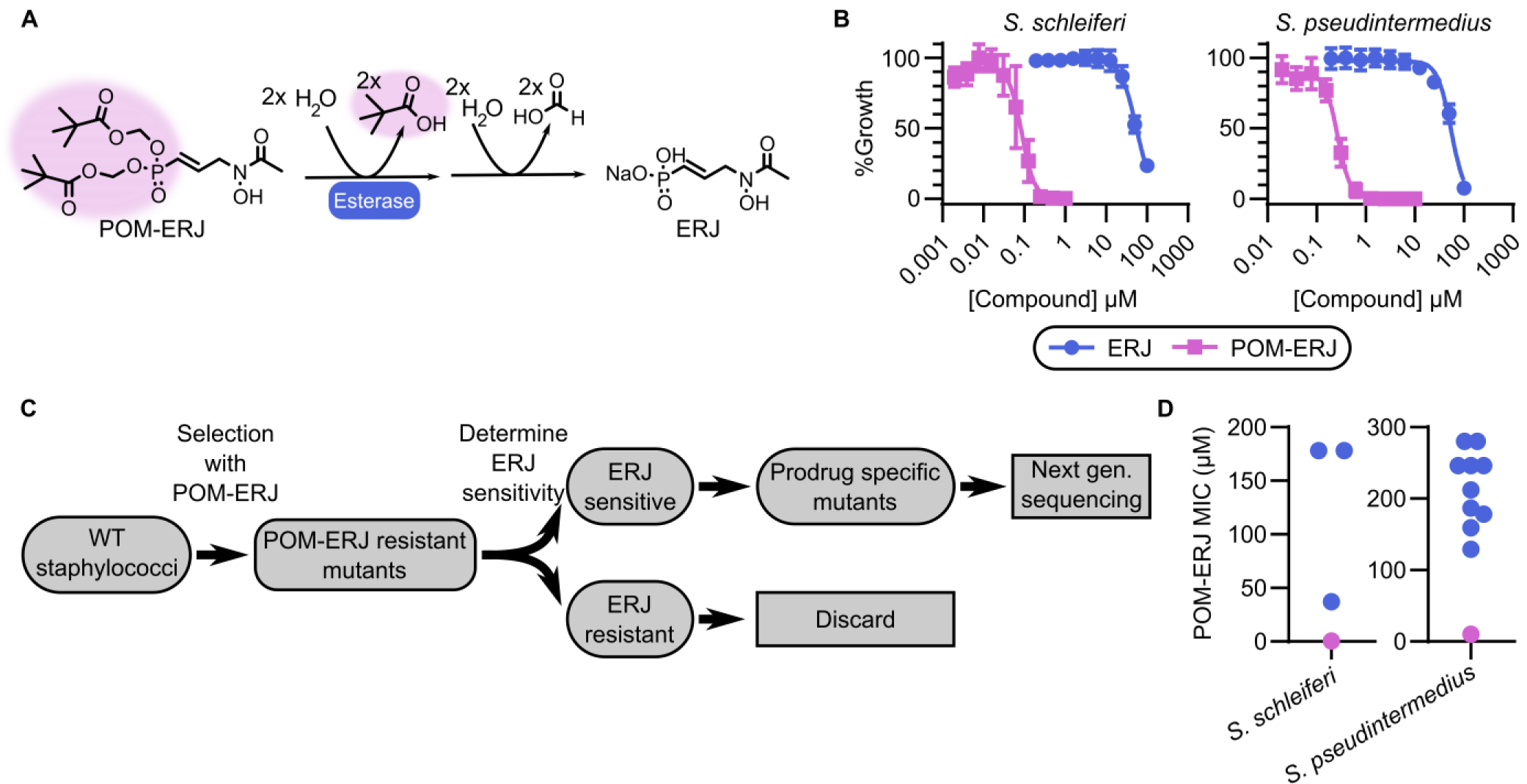
(A) Predicted POM-ERJ activation pathway. POM-promoiety highlighted in pink. (B) Dose-dependent growth inhibition of zoonotic staphylococci, *S. schleiferi* (left) and *S. pseudintermedius* (right), by ERJ (blue) and POM-ERJ (pink). Displayed values are the means ± SD of three independent experiments performed in technical duplicate. (C) Screening strategy to identify prodrug activating enzymes. (D) Distribution of MIC values for WT (pink) and POM-ERJ resistant mutants from *S. schleiferi* (left) and *S. pseudintermedius* (right). Displayed values are the means values for each strain from three independent experiments performed in technical duplicate.

In our strategy, we took advantage of inhibitor pairs with the same target engagement, with and without prodrug modification. We employed the phosphonate antibiotic ERJ, which selectively inhibits the intracellular enzyme deoxyxylulose phosphate reductoisomerase (DXR), and POM-ERJ, the bis-pivaloyloxymethyl prodrug form of ERJ, which inhibits intracellular DXR even though it has been shown to lack direct activity against purified recombinant DXR *in vitro* (27). We sought to enrich for staphylococcal strains that were resistant to prodrugged inhibitors (e.g. POM-ERJ) but remained sensitive to the parent phosphonate ERJ itself (27). For this reason, we first isolated staphylococcal colonies that arose from solid media containing POM-ERJ. Next, we screened these POM-ERJ-resistant isolates for cross-resistance to our parent compound, ERJ. POM-ERJ-resistant strains that remained sensitive to ERJ were subjected to whole genome sequencing to identify candidate genetic mutations giving rise to the resistance phenotype (Fig. 1C). To identify conserved resistance mechanisms, we performed this screen/counter-screen independently in two staphylococcal species, *S. schleiferi* and *S. pseudintermedius*. We isolated and characterized a total of 18 POM-ERJ-resistant staphylococcal strains, with MIC_90_ values ∼10-50 fold higher than that of the respective wild-type (WT) parental lines (Fig. 1D).

### POM-ERJ resistance does not alter cell wall size in staphylococci

In previous work, we and others have found that cellular entry of the phosphonate antibiotic ERJ and ERJ analogs requires the phosphonate transporter GlpT (16, 27, 34, 35). In contrast, entry of POM-ERJ is transporter-independent (16, 27). POM-ERJ resistance could therefore arise through cell wall modifications that directly disrupt cell penetration of prodrugs. Such cell wall alterations might therefore lead to cross-resistance to other antimicrobials, such as daptomycin or vancomycin. To establish the selectivity of POM-ERJ-resistance, we determined the antimicrobial sensitivity of a subset of our prodrug-resistant strains against a panel of 18 clinical antibiotics with diverse mechanisms-of-action. We find that POM-ERJ-resistant strains are not cross-resistant to other inhibitors, including daptomycin and vancomycin (Table S1), suggesting a prodrug-specific mechanism of resistance. Additionally, we quantified the cell wall size in POM-ERJ-resistant staphylococci by transmission electron microscopy, because an established daptomycin and vancomycin resistance strategy for *S. aureus* is the generation of thickened cell walls that reduce inhibitor entry (36, 37). We find no changes in cell wall thickness in prodrug-resistant isolates compared to their prodrug-sensitive WT parental lines (Fig 2).

**Figure 2.**
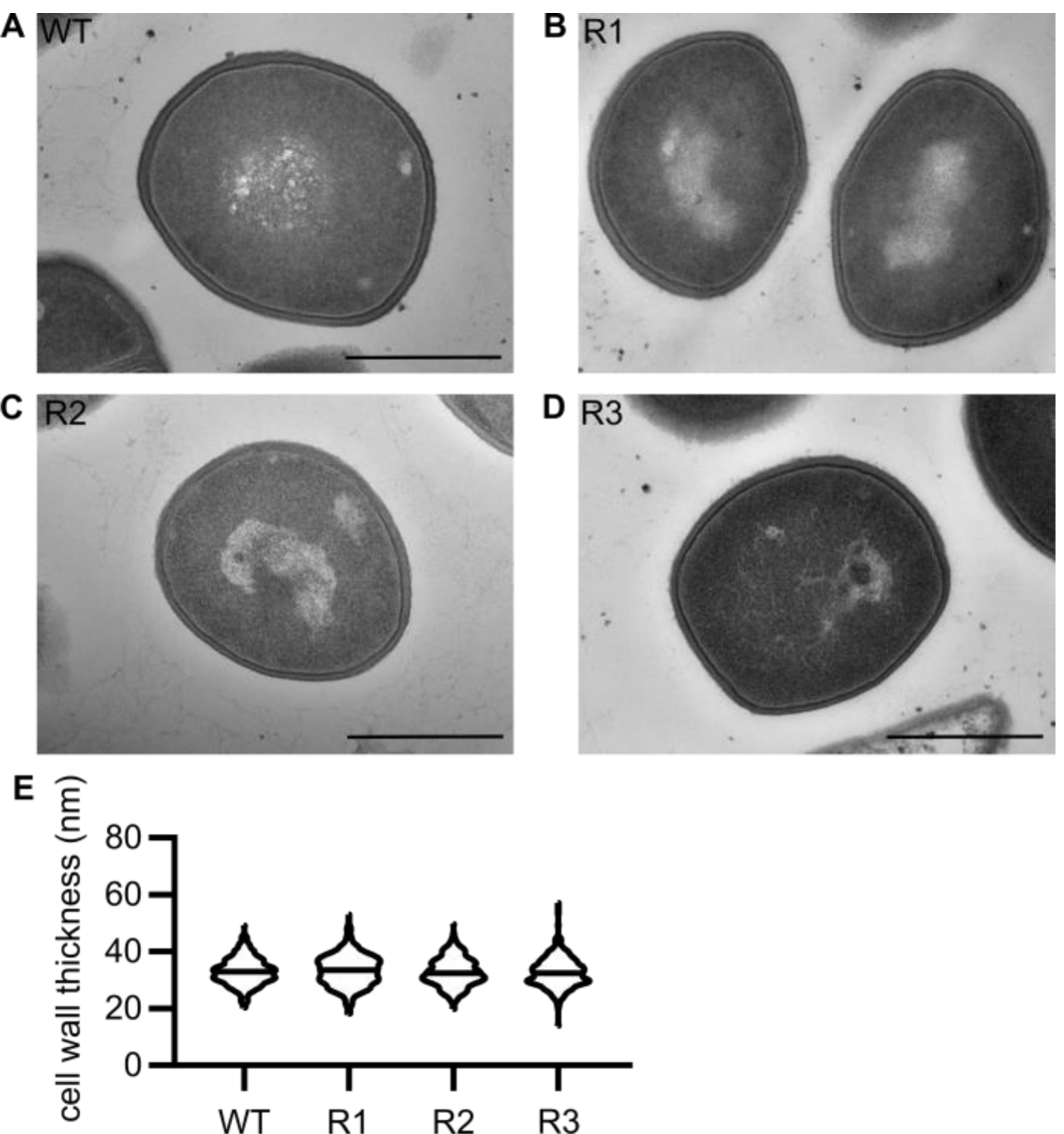
(A-D) Representative transmission electron micrographs of WT (A) or three independent POM-ERJ resistant *S. schleiferi* strains (B-D). Scale bars = 500 nm. (E) Distribution of cell wall thickness in WT and POM-ERJ resistant *S. schleiferi* as measured in a total of 300 cells from three independent experiments of 100 cells each. Midline indicates mean of all measurements.

### POM-ERJ-resistant staphylococci are cross-resistant to other carboxy ester prodrug antibiotics

If POM-ERJ resistance is due to loss of a prodrug activating enzyme(s), we hypothesized that POM-ERJ-resistant staphylococci would likewise be cross-resistant to other carboxy ester prodrug antibiotics. To evaluate this possibility, we selected several additional pairs of inhibitors (carboxy ester prodrugs and their cognate parent (non-prodrugged) compounds), with distinct cellular targets (e.g. penicillin binding protein, deoxyxylulose reductoisomerase (DXR), and enolase) (Fig. 3) (22, 32). For three of our POM-ERJ-resistant *S. schleiferi* isolates, we determined the minimum inhibitory concentration (MIC) for each compound, compared to the WT parental strain (Fig. 4).

**Figure 3.**
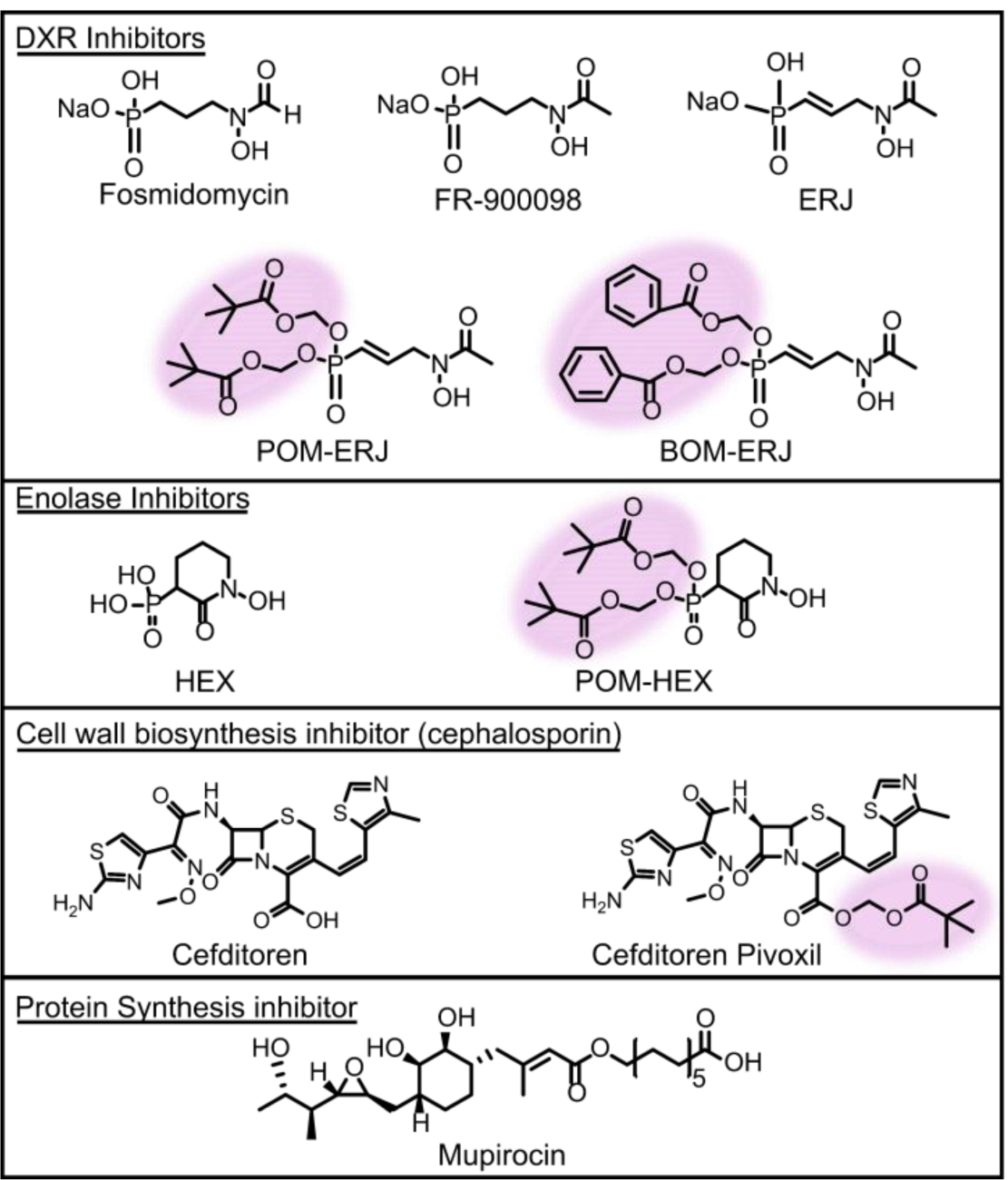
Structures of antistaphylococcal inhibitors used in this study. Structures are grouped by mechanism of action. For prodrugged compounds, promoieties are highlighted in pink.

**Figure 4.**
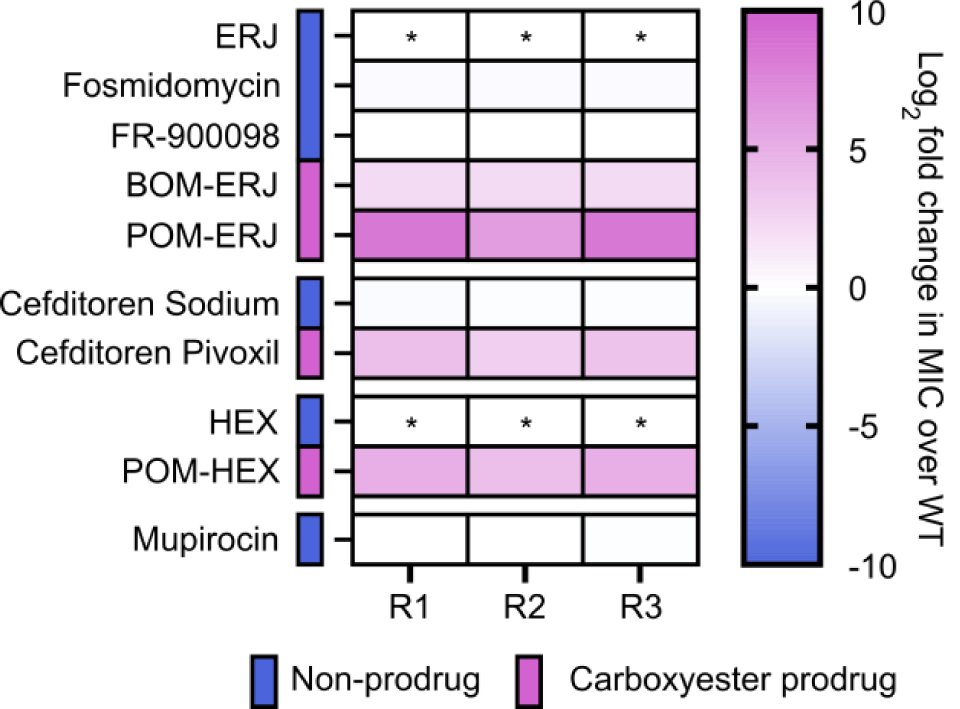
Cross-resistance to lipophilic ester prodrugs in POM-ERJ-resistant *S. schleiferi*. WT and POM-ERJ resistant *S. schleiferi* were treated with the compounds displayed in Figure 3. Compounds are grouped by mechanism of action and color coded to indicate whether a given compound is a carboxy ester prodrug. Displayed are the mean values of the fold change (resistant isolate/WT) of three independent experiments performed in technical duplicate. * indicates compounds whose MIC values were too high to measure. Numerical data additionally provided in Table S2.

We find that POM-ERJ-resistant staphylococci remain equally sensitive to non-prodrugged compounds (such as ERJ analogues) and the third-generation cephalosporin cefditoren. In contrast, POM-ERJ-resistant staphylococci exhibit significantly increased MICs to multiple classes of lipophilic ester prodrugs, exhibiting cross-resistance to both cefditoren pivoxil (cell wall inhibitor) and POM-HEX (inhibitor of enolase) (Fig. 4, Table S2). Thus, POM-ERJ-resistant staphylococci are cross-resistant to other POM-prodrug inhibitors, regardless of the intracellular target. Our data suggest that POM-prodrugs follow a common and conserved activation mechanism that has been disrupted in our POM-ERJ-resistant isolates.

To explore how changes in the chemical structure of the prodrug group impacts prodrug resistance, we also evaluated whether our POM-ERJ-resistant isolates were cross-resistant to antimicrobial prodrugs that possess another common carboxy ester prodrug moiety, benzyloxymethyl (BOM) (Fig. 3). Indeed, we find our POM-ERJ-resistant isolates are also cross-resistant to BOM-ERJ (Fig. 4).

Carboxy ester prodrugs are more lipophilic than their parental molecules. To evaluate whether prodrug resistance in our strains is driven by the lipophobicity of the molecule rather than its ester bond, we selected an additional highly lipophilic antibiotic, mupirocin, which inhibits protein biosynthesis (Fig. 3). POM-ERJ-resistant staphylococci were not cross-resistant to mupirocin, further supporting that prodrug resistance in these strains is specific to the carboxy ester bond of the prodrug (Fig. 4).

### POM-ERJ resistant staphylococci are enriched in mutations in the GloB gene

To characterize the genetic changes associated with carboxy ester prodrug resistance, we performed whole genome sequencing of prodrug resistant isolates of both *S. schleiferi* and *S. pseudintermedius*. The whole genomes of each isolate were compared to the respective parental genome and candidate genetic changes were verified by Sanger sequencing. We prioritized nonsynonymous genetic changes that were represented in more than one strain. A complete list of identified mutations is found in Table S3.

In both independent genetic screens, we found that prodrug resistant staphylococci were enriched in mutations in an evolutionarily conserved locus. We identified multiple isolates (3/16 *S. schleiferi*, 14/18 *S. pseudintermedius*) with sequence modifications in the locus annotated as hydroxyacylglutathione hydrolase, *gloB* (LH95_06060 in *S. schleiferi*, SPSE_1252 in *S. pseudintermedius*, Table S3). Most genetic changes in *gloB* were nonsynonymous single nucleotide polymorphisms, though two nonsense alleles that would truncate approximately 50% of the protein were also identified (Fig. 5, Table S3). In several strains, the only genetic variation that distinguished WT and resistant genomes was within the *gloB* locus.

**Figure 5.**
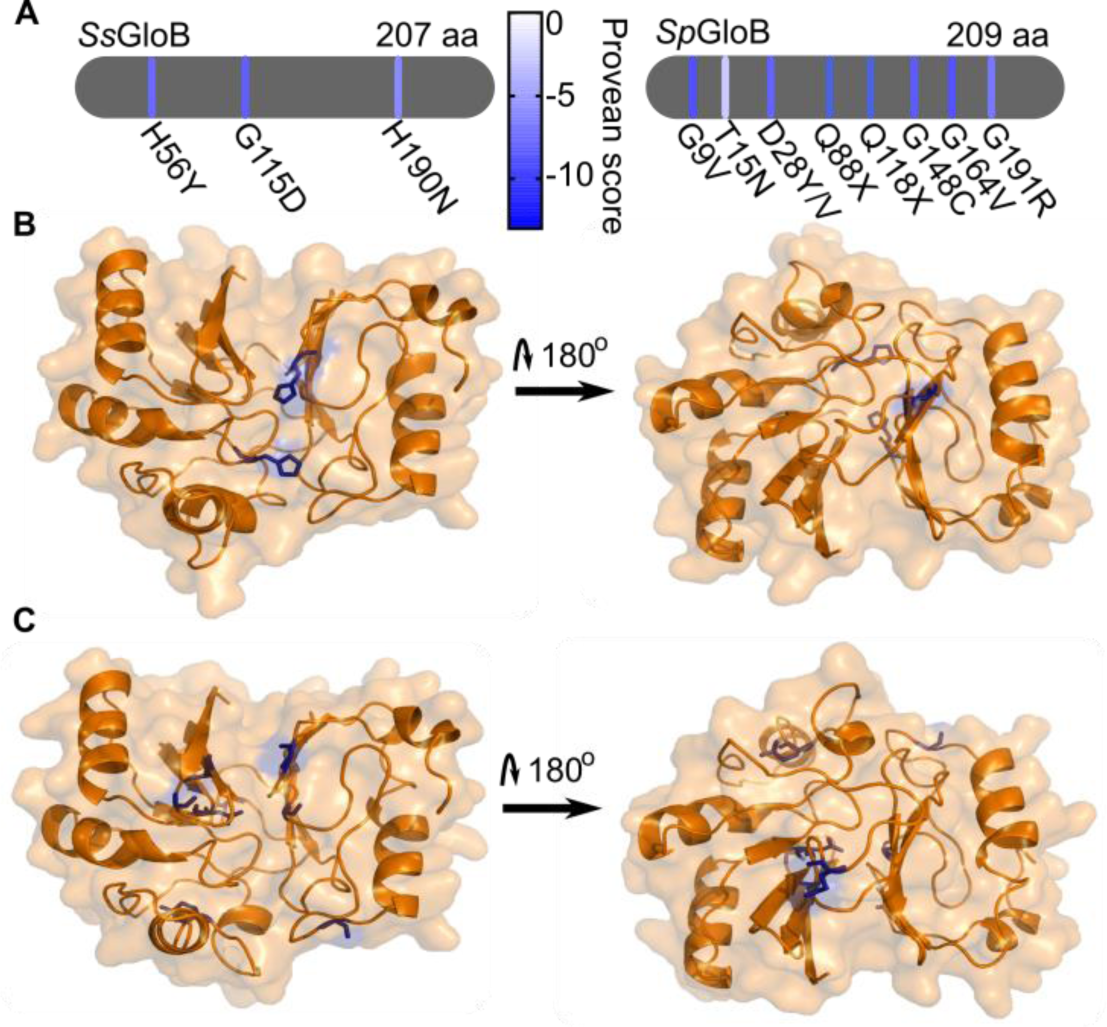
POM-ERJ resistant staphylococci are enriched for mutations in the locus encoding hydroxyacylglutathione hydrolase (GloB). (A) Locations and identities of GloB mutations discovered by whole-genome sequencing and independently verified by Sanger sequencing. Line coloring represents predicted impact of a given mutation on GloB function, scores below -2.5 are predicted to be deleterious. (B, C) Homology models of *S. schleiferi* (B) and *S. pseudintermedius* GloB generated using SWISS-MODEL. Residues found to be mutated in POM-ERJ resistant staphylococci explicitly shown in blue.

Of the 17 identified GloB mutations, 12 unique alleles were identified in prodrug-resistant staphylococci. Using PROVEAN, an algorithm which quantifies the predicted impact of amino acid substitutions on protein function, each of these 12 alleles is predicted to have deleterious effects on protein function (below the threshold score of -2.5) (Fig. 5) (38). *S. schleiferi* and *S. pseudintermedius* are non-model organisms that possess endogenous CRISPR-Cas9 systems and transformation of these organisms has not yet been described (39). Attempts to ectopically complement *gloB* mutant strains with WT GloB (>90 independent transformation attempts using established methods for *S. aureus, S. epidermidis*, and *B. subtilis*) were unsuccessful in recovering transformed colonies, despite preparing plasmid from the *S. aureus* restriction deficient cloning intermediate, RN4220, and the cytosine methyltransferase negative *E. coli* mutant, DC10B (40–46). However, the independent selection of 12 unique loss-of-function alleles in two different species strongly suggests that loss of GloB function is responsible for prodrug resistance in *S. schleiferi* and *S. pseudintermedius*.

### Structural basis of GloB loss-of-function

As prodrug-resistance mutations in GloB map along its entire linear sequence, we next examined the structural basis for GloB loss-of-function. We generated homology models of both *Ss*GloB and *Sp*GloB using SWISS-MODEL (47). The resulting staphylococcal model is based on the sequence-similar metallo-β-lactamase superfamily member from *Thermus thermophilus* (PDB 2ZWR) (48). This hit had a global model quality estimate (GNQE) of 0.71 and 0.70 for *S. schleiferi* and *S. pseudintermedius* GloB homologs, respectively, suggesting the built models are reliable and accurate. In both protein models, we find that POM-ERJ-resistance mutations are primarily located towards the interior of the protein, occupying the same cavity as the well conserved glyoxalase II metal binding motif (THxHxDH) (49). This modeling thus indicates that these prodrug-resistance alleles impair the GloB active site (Fig. 5).

### GloB is a functioning type II glyoxalase, not a β-lactamase

GloB is predicted to be a type II glyoxalase and a member of the large metallo-β-lactamase protein superfamily (INTERPRO IPR001279). Members of this superfamily hydrolyze thioester, sulfuric ester, and phosphodiester bonds, such as the ester linkage present in POM-ERJ (49– 52). Type II glyoxalases catalyze the second step in the glyoxalase pathway that is responsible for the conversion of methylglyoxal (a toxic byproduct endogenously produced during metabolism) to lactic acid. Specifically, GloB catalyzes the conversion of D-lactoylglutathione to D-lactate.

To determine whether *Ss*GloB encodes a functional type II glyoxalase, we evaluated whether *Ss*GloB hydrolyzes S-lactoylglutathione using an assay in which hydrolysis of S-lactoylglutathione is linked to a change in absorbance (Fig. 6A). We purified recombinant WT *Ss*GloB protein and its catalytically inactive variant, *Ss*GloB^H54N^, in which the histidine of the canonical metal binding motif (THxHxDH) has been altered to an asparagine (Fig. S1) (49, 51, 52). We find that *Ss*GloB, but not *Ss*GloB^H54N^, hydrolyzes S-lactoylglutathione with a specific activity of 0.493 μmol*min^-1^mg^-1^ (Fig. S2, Fig. 6B,C). This activity is similar to other characterized microbial type II glyoxalases (*Saccharomyces cerevisiae*, 1.34 μmol*min^-1^mg^-1^; *Trypanosoma brucei*, ∼8 μmol*min^-1^mg^-1^), but is much lower than that of previously characterized type II glyoxalases from plants and mammals (20-2000 μmol*min^-1^mg^-1^) (53–61). We determined the metal dependence of *Ss*GloB and find that *Ss*GloB is a functional type II glyoxalase in manganese, cobalt, calcium, and zinc, with a modest preference noted towards magnesium (Fig. S3).

**Figure 6.**
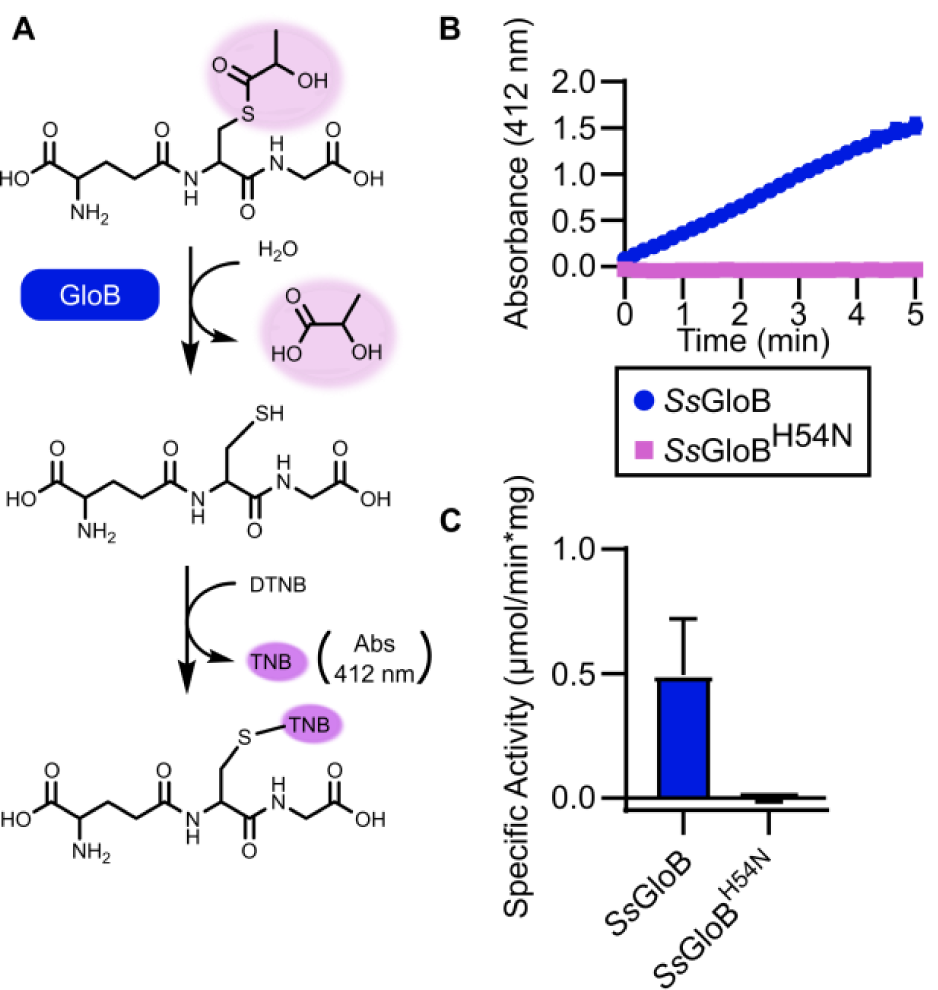
(A) Enzymatic catalysis of S-lactoylglutathione by GloB. DTNB conversion to TNB results in increased absorbance at 412 nm. (B) Reaction progress curve for *Ss*GloB (blue) and catalytically inactive *Ss*GloB H54N (pink), using S-lactoylglutathione as a substrate. (C) *Ss*GloB and *Ss*GloB H54N specific activity for S-lactoylglutathione. Displayed are the means ± SD from three independent experiments preformed in technical duplicate.

As some members of the metallo-β-lactamase protein superfamily mediate hydrolysis of β-lactam antibiotics, we considered whether GloB also had β-lactamase activity. Because *gloB* mutant strains are not cross-resistant to the β-lactam-containing antibiotics (except for the prodrugged cephalosporin, cefditoren pivoxil) (Fig. 4, Table S2), we predicted that GloB was not a functional metallo-β-lactamase. As expected, we find that *Ss*GloB does not hydrolyze the β-lactamase ring of nitrocefin (a canonical β-lactamase substrate), in contrast to the active *B. cereus* β-lactamase (Fig. S2).

### Staphylococcal GloB hydrolyzes POM-ERJ *in vitro* and *in vivo*

Loss-of-function mutation in GloB is associated with resistance not only to POM-ERJ, but also to other ester prodrugs. Because GloB does not mediate resistance to ERJ or other phosphonates, our data suggested that GloB might directly catalyze the conversion of POM-ERJ to ERJ. To determine whether GloB de-esterifies POM-ERJ, we developed a liquid chromatography-mass spectrometry (LC-MS)-based assay to quantify POM-ERJ concentrations. Incubation of purified recombinant *Ss*GloB protein, but not its inactive variant (*Ss*GloB^H54N^), with POM-ERJ results in rapid loss of POM-ERJ, consistent with *Ss*GloB-mediated cleavage (Fig. 7A). To determine whether prodrug activation activity is conserved among staphylococcal GloB homologs, we also purified recombinant GloB from the human pathogen *S. aureus* (Fig. S1). We find that *Sa*GloB also directly hydrolyzes POM-ERJ (Fig. 7A).

**Figure 7.**
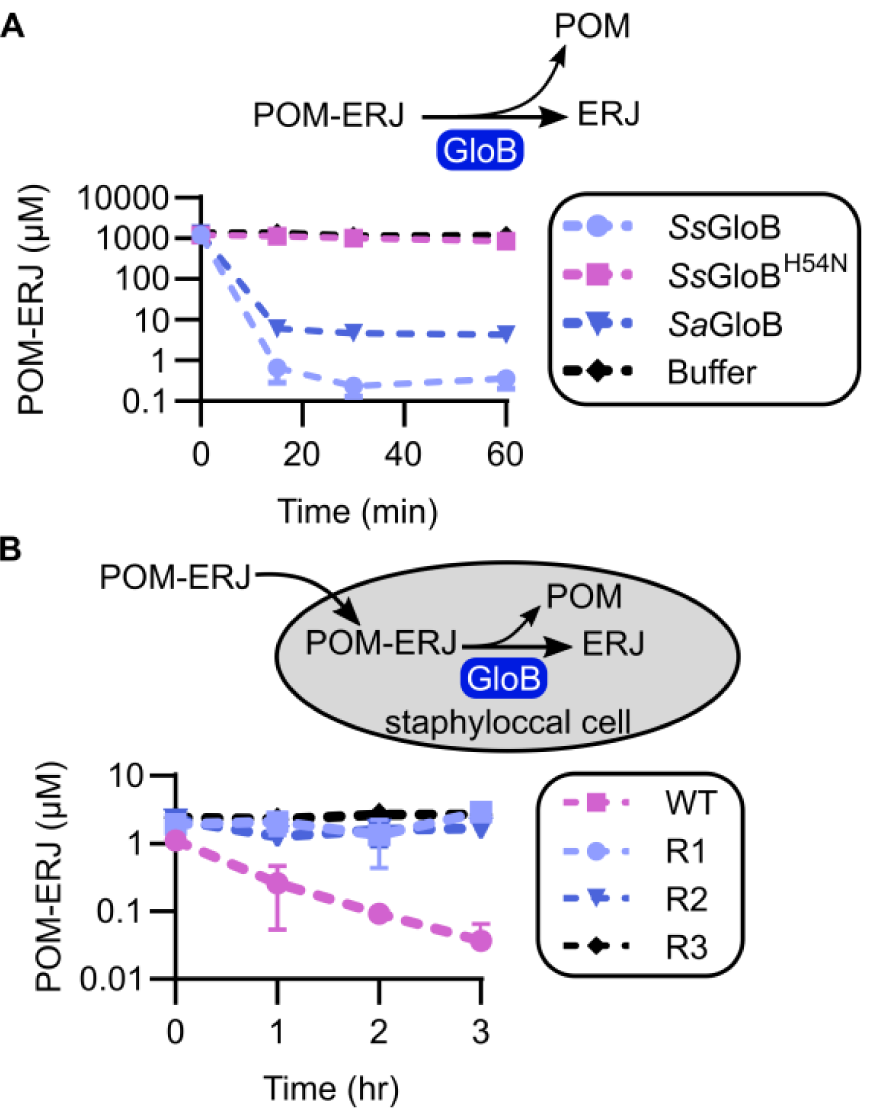
(A) Recombinant *Ss*GloB, catalytically inactive *Ss*GloB H54N, GloB from *S. aureus* (*Sa*GloB), or buffer were incubated with POM-ERJ and prodrug concentrations were measured by LC-MS. (B) Wild-type and POM-ERJ-resistant *gloB* mutant *S. schleiferi* isolates were treated with POM-ERJ and intracellular drug concentrations were measured by LC-MS. Displayed are the mean values ± SD from three independent experiments. Error bars may not be visible due to precision in measurement.

To determine whether GloB mediates intracellular prodrug activation, we evaluated the intracellular concentrations of POM-ERJ in drug-treated WT and *gloB* mutant staphylococci. We prepared staphylococcal cultures treated with POM-ERJ and quenched the reaction at several timepoints to monitor the course of intracellular prodrug depletion. As expected, we find that POM-ERJ is rapidly depleted in WT *S. schleiferi*, consistent with enzymatic activation. In contrast, POM-ERJ concentrations do not decrease over time in *gloB* mutant strains, in which the sole genetic change in each strain compared to WT is in the *gloB* locus (Fig. 7B). This suggests that the initial step in carboxy ester prodrug activation in staphylococci lacks functional redundancy and is exclusively dependent on GloB.

### Staphylococcal GloB enzymes represent a distinct clade of bacterial glyoxalases

Because staphylococcal GloB mediates de-esterification of ester prodrugs, we sought to evaluate the feasibility of using these enzymes to design prodrugs specifically targeted for activation in staphylococci. We constructed a phylogenetic tree of GloB homologs across diverse microbial genomes, as well as in humans and mice (Fig. S4A), specifically including sequences of previously characterized GloB homologs. We find that considerable sequence variation exists within GloB homologs, with no clear clustering by phylogeny except for those GloB homologs originating in plants and mammals. This contrasts with a phylogenetic tree generated using the DNA-directed RNA polymerase subunit beta (rpoB), which generally follows the traditional tree of life (Fig. S4B).

While sequence differences between staphylococcal GloB and human GloB suggest that there may be substrate utilization differences between humans and staphylococci, ultimately differences within the active site are likely to drive substrate specificity. Using pymol, we aligned our homology model of *Ss*GloB with the glutathione bound GloB from humans (PDB ID: 1qh5) (62, 63). The two structures align well with a root-mean-square deviation (RMSD) of 1.528Å, and are well conserved in the overall structure as well as the characteristic Zn binding motif, THxHxDH (Fig. S5A,B). Notably, however, *Hs*GloB has a significant C-terminal extension which is not present in *Ss*GloB. This C-terminal extension forms an α-helix which borders the active site and contains two residues, K252 and R249, which appear to be involved in coordinating the co-crystallized glutathione substrate (Fig. S5C). The absence of this C-terminal extension in our *Ss*GloB homology model suggests that *Hs*GloB and *Ss*GloB have distinct active site chemistry that may be exploited to drive prodrug activation selectively by *Ss*GloB vs *Hs*GloB.

## DISCUSSION

Antimicrobial resistance is a substantial challenge for treatment of both human and animal staphylococcal infections. Widespread methicillin resistance contributes both to poor clinical outcomes and increased treatment costs, and resistance is emerging to agents of last resort such as vancomycin and linezolid (7). Current antimicrobial therapies target a fraction of essential cellular processes, and metabolism remains a promising area for therapeutic development (11, 12). Many metabolic genes are essential for growth, especially in the nutrient limited setting of infection (64–67). Additionally, chemical ligands are readily designed with high potency by mimicking natural substrates used by metabolic enzymes. Finally, because active site mutations that disrupt binding of competitive inhibitors are likely to deleteriously affect enzyme function, the barrier to resistance can be high (68, 69). Although many metabolic processes are conserved between humans and microbes, selective targeting of microbes is achievable as is demonstrated by the success of folate antagonists (trimethoprim/sulfamethoxazole) and bedaquiline (a F_0_F_1_ ATP synthase inhibitor of *Mycobacterium tuberculosis*) (70–72).

Unfortunately, many metabolic inhibitors require cell-impermeable phosphonic acids for efficient target inhibition. Prodrugging strategies to increase cellular penetration have been developed for a variety of therapeutics, most notably the anti-cancer and anti-viral nucleosides (19). These prodrug strategies must be labile-enough that the compound is activated within the target cell, yet stable enough to resist premature prodrug activation by the sera. Prodrugs which are selectively activated within target cells have the added benefit of reducing off-target toxicity effects. To achieve cell-targeted prodrug activation, knowledge of the activation mechanisms in sera, as well the target cell, are essential. While prodrug targeting has been achieved for liver therapies, this strategy has yet to be employed for bacterial antibiotics that employ ester prodrug moieties (33).

In this work, we have identified a new mechanism for the de-esterification and activation of lipophilic ester prodrugs though a conserved staphylococcal esterase in the metallo-β-lactamase superfamily. Loss-of-function of GloB confers resistance to lipophilic carboxy ester prodrugs in two zoonotic pathogens, *S. schleiferi* and *S. pseudintermedius* (Fig. 1D, Table S3). Purified recombinant GloB from *S. schleiferi* and the related human pathogen *S. aureus* directly catalyzes pro-drug de-esterification *in vitro* (Fig. 7A). Because *gloB* mutant staphylococci are cross-resistant to other POM-containing prodrugs that differ in “warhead” and intracellular targets (Fig. 4), we propose that substrate-specificity of GloB appears driven by recognition of the lipophilic promoiety, rather than the target inhibitory portion of each compound.

Bacterial prodrug ester activation through GloB hijacks a conserved bacterial protective mechanism in bacteria, as hydroxyacylglutathione hydrolase represents the second enzyme of the two-step glyoxalase pathway. During normal metabolism, the glycolytic intermediates glyceraldehyde-3-phosphate (GAP) and dihydroxyacetone phosphate (DHAP) undergo nonenzymatic decomposition to methylglyoxal, a toxic metabolite. GloB is required for methylglyoxal detoxification, as methylglyoxal is highly reactive and irreversibly glycates proteins and nucleic acids (73–75). In *S. aureus*, methylglyoxal accumulation potentiates antibiotic susceptibility (76). In addition, methylglyoxal is itself directly antibacterial and postulated to be the primary antistaphylococcal ingredient in Manduka honey (used on chronic wounds) (76–79). Our studies suggest that strains of *S. schleiferi* and *S. pseudintermedius* lacking GloB have preserved axenic growth in rich media, which raises concern for the ease of resistance development when GloB-targeted prodrugs are used as anti-infectives. However, the known toxicity of methylglyoxal in a host infection setting suggests that reduced methylglyoxal detoxification as the result of GloB loss-of-function would not be well tolerated *in vivo*.

Identification of GloB as a prodrug activating enzyme in staphylococci is a major step forward for highly selective microbial targeting of compounds. Though GloB homologs are widespread in microbes and are present in humans, significant sequence variation exists in GloB sequences, which results in a variety of GloB substrate preferences (Fig. S4). For example, human GloB has an additional α-helix along the active site that introduces two additional residues, K252 and R249 to the substrate binding pocket (Fig. S5) (63). These residues, and this α-helix, are notably absent in microbial GloBs, suggesting that there are underlying substrate differences between human and microbial GloB enzymes. Furthermore, there is substantial sequence variation in GloB orthologs across all microbes, suggesting that GloB substrate specificities may discern between individual clades of bacteria. We expect that development of prodrugs specific to GloB would result in a narrow-spectrum antibiotic which would reduce off-target effects on the microbiome and decrease the broad pressure to evolve resistance.

## Materials and Methods

### Inhibitors

Fosmidomycin (Millipore Sigma) and FR-900098 (Millipore Sigma) were resuspended in sterile water. POM-ERJ and POM-HEX were synthesized and stored in DMSO as described (29, 32). Cefditoren pivoxil (Millipore Sigma), cefditoren sodium (Clearsynth), and mupirocin (Millipore Sigma) were resuspended in DMSO. The synthesis of BOM-ERJ will be reported elsewhere.

### Generation of POM-ERJ-resistant mutants in *S. schleiferi* and *S. pseudintermedius*

Clinical isolates of *S. schleiferi* (S53022327s) and *S. pseudintermedius* (H20421242p) were cloned and adapted to laboratory media through three rounds of sequential colony isolation and growth on Luria Broth (LB) agar plates. The isolated POM-ERJ-sensitive parental clones were incubated overnight on LB agar containing POM-ERJ at 3.56 µM and 7.12 µM for *S. schleiferi* and 11.2 µM and 22.4 µM for *S. pseudintermedius*. Surviving single colonies were re-struck onto LB agar for clonal isolation. POM-ERJ resistance of isolated clones was confirmed by overnight growth on LB agar containing POM-ERJ (3.56-22.4 µM). The POM-ERJ-sensitive parental clones were used as a control to confirm growth and antibiotic resistance.

### Quantification of resistance

Minimum Inhibitory Concentration (MIC) assays were performed using microtiter broth dilution in clear 96-well plates (80). Compounds were serially diluted in duplicate for a total of 10 serial dilutions. Top well concentrations were: POM-ERJ 280 µM, BOM-ERJ 53.95 µM, KMH-102 53.95 µM, cefditoren pivoxil 201.38 µM, cefditoren sodium 56.65 µM, POM-HEX 100 µM, mupirocin 2.50 µM, FR-900098 1 mM, fosmidomycin 100 µM. Bacteria cultured without drug were used as a positive control for growth, and LB without bacteria was used as a negative control for contamination. Plates were inoculated with 75 µL bacteria diluted to 1 × 10^5^ CFU/mL in LB. After inoculation, plates were incubated for 16-24 h while shaking at 200 RPM at 37°C. Plates were visually inspected, and the lowest concentration of antibiotic suppressing visual growth was recorded as the MIC. All experiments were performed at least in triplicate and data reported represent the mean ± SD.

### Transmission Electron Microscopy

For ultrastructural analysis, bacteria were cultured in 5 mL LB while shaking at 37°C until OD_600_ = 0.25-1.0. A 1 mL sample of exponential phase bacteria was pelleted at 6,000 rcf and resuspended in 1 mL fix (2% paraformaldehyde/2.5% glutaraldehyde (Polysciences Inc., Warrington, PA) in 100 mM sodium cacodylate buffer, pH 7.2) for 1 h while rocking at RT. The fixed suspension of bacteria was washed in sodium cacodylate buffer and postfixed in 1% osmium tetroxide (Polysciences Inc.) for 1 h. Samples were then rinsed extensively in dH_2_O prior to en bloc staining with 1% aqueous uranyl acetate (Ted Pella Inc., Redding, CA) for 1 h. Following several rinses in dH_2_O, samples were dehydrated in a graded series of ethanol and embedded in Eponate 12 resin (Ted Pella Inc.). Sections of 95 nm were cut with a Leica Ultracut UC7 ultramicrotome (Leica Microsystems Inc., Bannockburn, IL), and stained with uranyl acetate and lead citrate. Samples were viewed at 30,000X on a JEOL 1200EX transmission electron microscope (JEOL USA, Peabody, MA) equipped with an AMT 8 megapixel digital camera (Advanced Microscopy Techniques, Woburn, MA). Cell wall thickness was measured (ImageJ 1.38g customized for AMT images) for 100 bacteria in three independent samples (total n = 300).

### Whole genome sequencing and variant discovery

Using a standard phenol-chloroform extraction and ethanol precipitation protocol, genomic DNA was isolated from overnight cultures of *S. pseudintermedius* and *S. schleiferi*. Sequencing libraries were prepared and sequenced by the Washington University Genome Technology Access Center (GTAC). 1 µg of DNA was sonicated to an average size of 175 bp. Fragments were blunt ended and had an A base added to the 3’ end. Sequence adapters were ligated to the ends and the sequence tags were added via amplification. Resulting libraries were sequenced on an Illumina HiSeq 2500 to generate 101 bp paired end reads. DNA quantity and quality were assessed by GTAC using Agilent Tapestation.

For the analysis, sequences from GenBank were retrieved from the following organisms: *S. pseudintermedius* ED99 (accession number CP002478) and *S. schleiferi* 1360-13 (CP009470) assemblies were downloaded from NCBI (ftp://ftp.ncbi.nlm.nih.gov). Paired-end reads were aligned to each of the available genomes using Novoalign v3.03. (Novocraft Technologies). Duplicates were removed and variants were called using SAMtools (81). SNPs were filtered against parent variants and by both mean depth value and quality score (minDP =5, minQ = 30) (82). Genetic variants were annotated using SnpEff v4.3 (83). For all samples, at least 90% of the genome was sequenced at 20x coverage. Whole genome sequencing data is available in the NCBI BioProject database and Sequence Read Archive under the BioProject ID 648133.

### Sanger sequencing of *S. schleiferi* and *S. pseudintermedius* variants

The SNPs, the reference sequences, and gene specific primers can be found in Table S4 for both *S. schleiferi* and *S. pseudintermedius*. Amplicons were sequenced by GENEWIZ.

### Staphylococcal GloB homology modelling

SWISS-MODEL (https://swissmodel.expasy.org/) was used to generate homology models. Modeling parameters were left at default. Both *Ss*GloB and *Sp*GloB models were built using the solved Metallo-β-lactamase superfamily protein, 2ZWR.1.A, which is 39.2% identical in sequence.

### Recombinant expression and purification of GloB

WT GloB from *S. schleiferi* was amplified using the forward and reverse primers in Table S4. The PCR product was then cloned into the BG1861 vector by ligation-independent cloning to introduce a N-terminal 6xHis tag and transformed into Stellar™ chemically competent cells (Clontech Laboratories) for plasmid propagation (84). Proper insertion was verified using restriction digest and Sanger sequencing. For *S. schleiferi* protein expression, the plasmid was transformed into *E. coli* Arctic Express (Agilent). Cells were grown to OD_600_ = 0.4-0.7, chilled to 8°C, and GloB expression was induced with 0.5 mM isopropyl-β-D-thiogalactoside (IPTG) overnight. For *S. aureus* protein expression, the plasmid was transformed into *E. coli* BL21 (DE3) pLysS cells (Promega). Cells were grown to OD_600_ = 0.4-0.7 and GloB expression was induced with 0.5 mM IPTG for 2 h. Cells were harvested by centrifugation at 4274 x g for 5 min at 4°C. The cell pellet was lysed by sonication in 50 mL lysis buffer containing 25 mM Tris HCl (pH 7.5), 20 mM imidazole, 1 mM MgCl_2_, 1 mM dithiothreitol (DTT), 1 mg/mL lysozyme, 75 U benzonase and 1 Complete Mini EDTA-free protease inhibitor tablet (Roche Applied Science). Insoluble proteins were removed by centrifugation twice at 20,000 x g for 20 min each. The hexahistidine-tagged DXR protein was affinity purified from soluble lysate via nickel agarose beads (Gold Biotechnology). Bound protein was eluted in 300 mM imidazole, 25 mM Tris HCl (pH 7.5), 1 mM MgCl_2_, 10% glycerol, and 250 mM NaCl. Affinity purified protein was further purified over a HiLoad 16/60 Superdex 200 gel filtration column (GE Healthsciences) using an AKTAExplorer 100 FPLC (GE Healthsciences). FPLC buffer contained 25 mM Tris HCl (pH 7.5), 250 mM NaCl, 1 mM MgCl_2_ and 10% glycerol. Fractions containing >90% pure enzyme (evaluated by SDS-PAGE) were concentrated by centrifugation using Amicon Ultra-15 centrifugal filter units (EMD Millipore) and flash frozen in liquid nitrogen before permanent storage at -80°C. Protein identity was verified using mass spectrometry at the University of Nebraska.

### GloB mutant generation

WT GloB for *S. schleiferi* was synthesized by GeneWiz, Inc (Beijing, China) with a CAT->AAT mutation in the 54th codon (H54N) and cloned into the BG1861 vector to introduce an N-terminal 6xHis tag. Proper insertion was verified by Sanger sequencing.

### Glyoxalase II activity assay

*S. schleiferi* GloB was tested for type II Glyoxalase activity as previously with minor changes (50). 50 μL reactions containing 25 mM Tris pH 7.5, 250 mM NaCl, 1 mM divalent salt, 10% glycerol, 200 μM 5,5′-Dithiobis(2-nitrobenzoic acid) (DTNB, Sigma D8130), and 1 mM D-lactoylglutathione (Sigma L7140) were monitored in a 96-well plate for an increase in absorbance at 412 nm. Reactions were pre-incubated at 37°C and initiated with the addition of GloB. The conversion of DTNB to the yellow colored substrate, TNB, by glutathione produced by GloB, was measured through time at 37°C and 412 nm. Assays were carried out over a range of GloB concentrations to ensure that the reaction rates are linear over the period of the assay. Assays were performed using Zn^+2^, Mn^+2^, Mg^+2^, Co^+2^, and Ca^+2^.

### Sample preparation for GloB vs. POM-ERJ mass spectrometry analysis

Reactions containing 25 mM Tris HCl (pH 7.5), 250 mM NaCl, 10% glycerol, 1 mM MnCl_2_, and 1 mM POM-ERJ were pre-warmed to 37°C before addition of WT GloB, catalytically inactive GloB (H54N), boiled GloB, or an equal amount of protein storage buffer to a final concentration of 1 μM. Reactions were placed at 37°C and sampled at 0, 15, 30, 60, 90, and 120 min. A 50 µL sample was withdrawn from each reaction at the times indicated, and the sample reaction was quenched by the addition of 200 µL acetonitrile containing 100 ng/μL enalapril as an internal standard. The samples were immediately frozen on dry ice and stored at -80°C until analysis.

The quenched reaction mixtures were centrifuged at 3200 rpm for 5 min, and 2 μL of the supernatant was diluted to 500 µL with water containing 100 ng/mL enalapril as an internal standard. Samples were analyzed by LC-MS/MS using an Applied Biosystems-Sciex API 4000. Analyte/internal standard peak area ratios were used to determine concentration and evaluate stability. Standards were evaluated over the range of 1 ng/mL to 1000 ng/mL. The MRM transitions for enalapril and POM-ERJ were m/z: 376.9 > 91.2 and 424.0 > 364.0, respectively. A Phenomenex Luna Omega polar C18 column (2.1 × 50 mm, 5 μm) was used for chromatographic separation. Mobile phases were 0.1% formic acid in water and acetonitrile with a flow rate of 0.5 mL/min. The starting phase was 1% acetonitrile increased to 100% acetonitrile over 0.9 min. Peak areas were integrated using Analyst Software (AB Sciex, Foster City, CA).

### In vivo cleavage of POM-ERJ

*S. schleiferi* cultures of WT and POM-ERJ^R^ strains were grown to an OD_600_ = 0.5-0.8 and then treated with 1 μM of POM-ERJ. The cultures were grown shaking at 37°C and 200 rpm and 50 μL were sampled at 0, 1, 2, and 3 h. The reactions were quenched by pelleting the cells at 4274 x g at 4°C and resuspending in 200 μL of acetonitrile with 100 ng/μL enalapril as an internal standard. The reactions were repeated in triplicate for each timepoint and strain. The LC-MS analysis was performed as described above.

## Acknowledgments

The authors are grateful to Joe Jez for ongoing support and helpful discussions. Financial support provided by NIH AI123433 to CSD and the GWU Department of Chemistry. A.O.J. is supported by NIH/NIAID R01-AI103280, R21-AI123808, and R21-AI130584, and AOJ is an Investigator in the Pathogenesis of Infectious Diseases (PATH) of the Burroughs Wellcome Fund.

## Notes

### Competing Interest Statement

The authors (AROJ and CSD) declare their status as co-inventors of U.S. provisional patent 62/686,416 filed June 18, 2018.

