## Supplemental methods, figures, and tables for "Antimicrobial prodrug activation by the staphylococcal glyoxalase GloB"

### Supplementary Methods

**β-lactamase activity assay.** *S. schleiferi* GloB was tested for β-lactamase activity using the chromogenic cephalosporin substrate Nitrocefin (Sigma Aldrich 484400) as in (85) but with minor changes. 50 μL reactions containing 25 mM Tris HCl (pH 7.5), 250 mM NaCl, 1 mM $MgCl_2$, 10% glycerol, and 200 μM Nitrocefin were preincubated for 15 min at 37$^oC$, and reactions were initiated upon addition of GloB. Cleavage of Nitrocefin was allowed to proceed at 37°C and tracked kinetically at 486 nm. Assays were carried out over a range of GloB concentrations starting at 2 g of protein (1.6 M).

**Phylogenetic tree construction.** The sequences of *S. schleiferi* GloB and RpoB homologs were retrieved from NCBI using BlastP against each specified organism. Organisms were selected to represent a wide array of commensal and pathogenic bacteria (86). Additional sequences were added from *Mus musculus*, *Homo sapiens*, and other previously characterized GloB orthologs for additional comparison. Sequence alignment was performed using MUSCLE, and visualized using iTOL (87, 88)

### Supplementary Figures

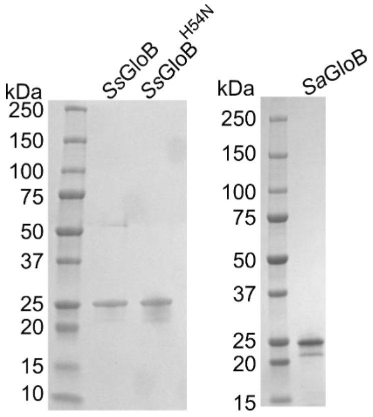

Figure S1. SDS-PAGE/Coomassie of purified recombinant *Ss*GloB, *Ss*GloB$^{H54N}$, and *Sa*GloB.

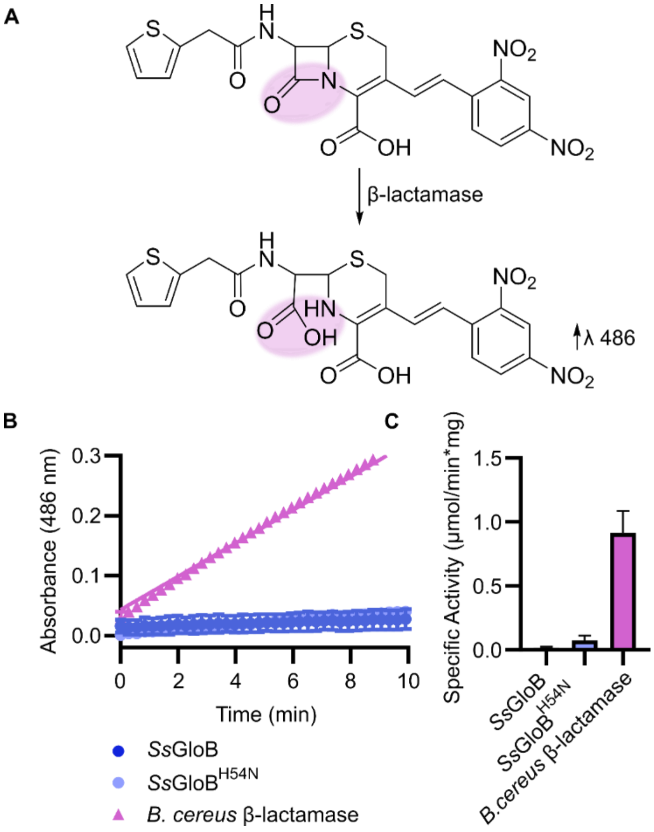

Figure S2. (A) Nitrocefin activation mechanism. Cleavage of the β-lactam ring results in increased absorbance at 486 nm. (B) Progress curve for nitrocefin cleavage by *Ss*GloB, *Ss*GloB$^{H54N}$, and commercially available β-lactamase from *B. cereus*. (C). Specific activity for *Ss*GloB, *Ss*GloB$^{H54N}$, and *B. cereus* β-lactamase against nitrocefin.

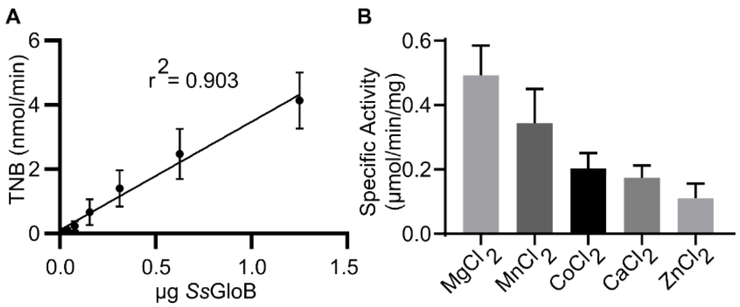

Figure S3. Assay validation for *Ss*GloB S-lactoylglutathione cleavage and detection via DTNB. (A) S-lactoylglutathione cleavage rate as a function of increasing *Ss*GloB. (B) *Ss*GloB metal dependence for cleavage of S-lactoylglutathione. *Ss*GloB was incubated with 1 mM of each divalent salt prior to S-lactoylglutathione reaction initiation.

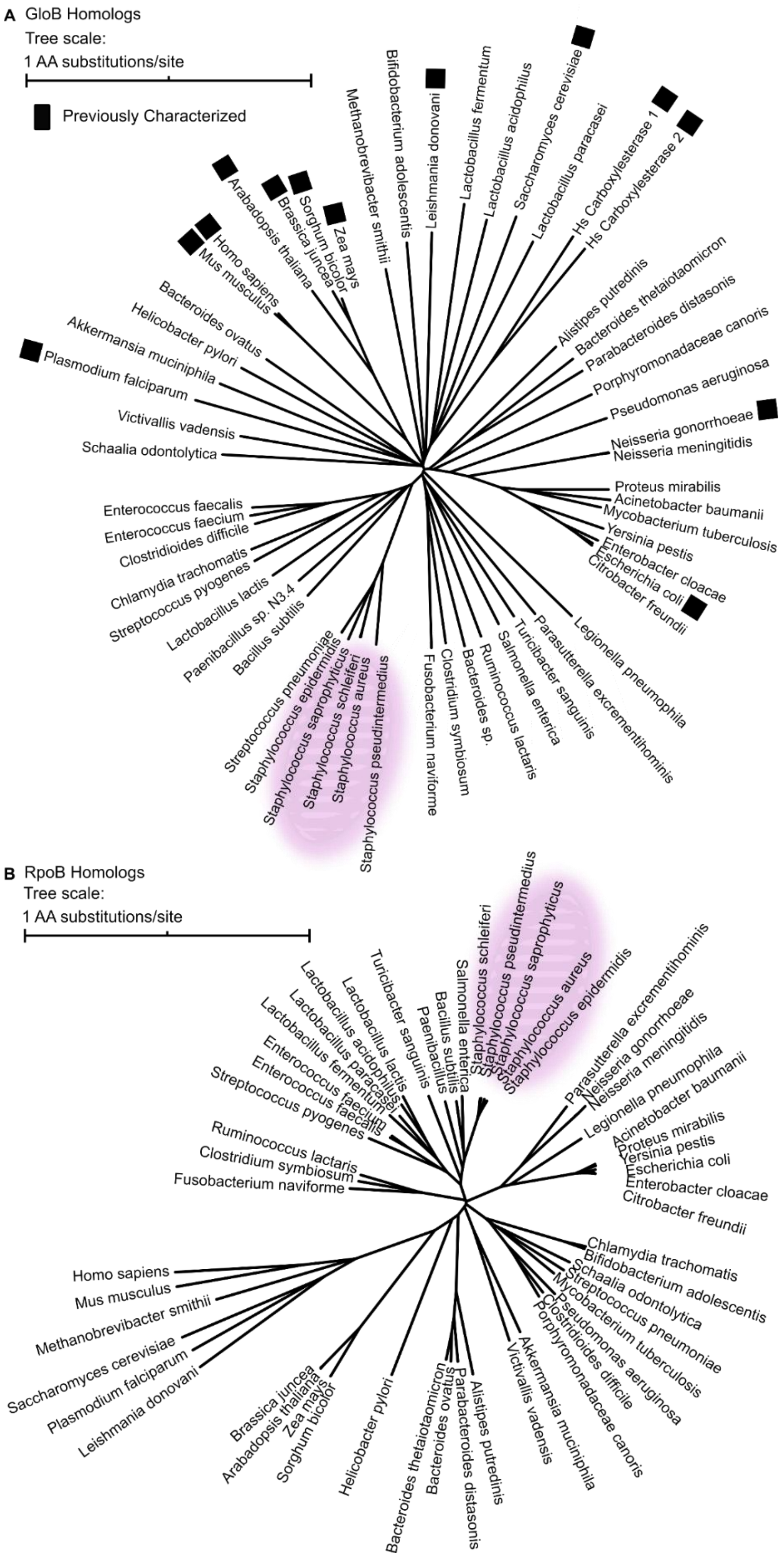

Figure S4. Phylogenetic trees of GloB (A) and RpoB (B) sequences. Pink highlighting overlain on staphylococcal species.

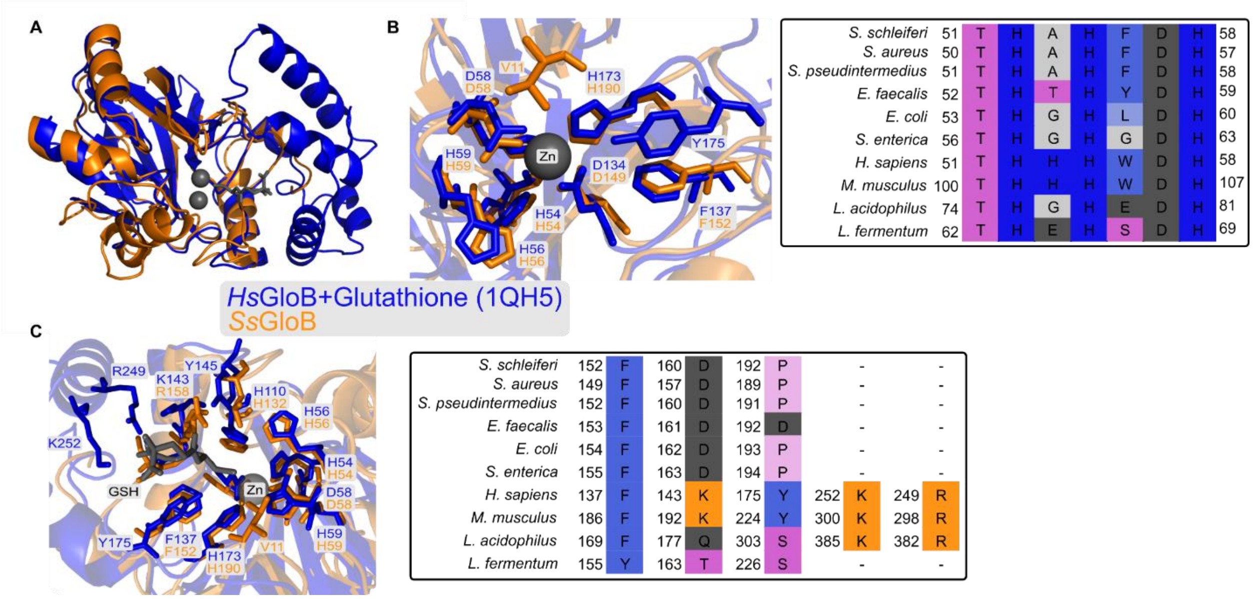

Figure S5. Alignment between *Hs*GloB (orange, PDB ID: 1qh5) and *Ss*GloB (blue) homology model. (A) Overall protein alignment, RMSD = 1.528Å. (B) Metal binding pocket (left), sequence alignment of residues contacting the bound Zn (*Hs*GloB) and their analogous residues colored according to amino acid chemical properties (right). (C) Substrate binding pocket (left), sequence alignment of residues contacting the bound glutathione colored according to amino acid chemical properties (GSH, right).

**Supplemental Tables**

Table S1: Zones of inhibition for POM-ERJ resistant zoonotic staphylococci against common frontline therapeutics. Presented are the zones of inhibition and whether the isolate is sensitive to the therapeutic (S), resistant (R), or intermediate (*) according to Clinical and Laboratory Standards Institute (CLSI) standard breakpoints.

| | | **WT S. pseudintermedius** | | **Sp80xa** | | **Sp80xb** | | **Sp80xc** | | **Sp80xd** | |
|---|---|---|---|---|---|---|---|---|---|---|---|
| **Antimicrobial** | **Mechanism of Action** | **Zone (in mm)** | **S/R?** | **Zone (in mm)** | **S/R?** | **Zone (in mm)** | **S/R?** | **Zone (in mm)** | **S/R?** | **Zone (in mm)** | **S/R?** |
| Cefoxitin | Cell wall synthesis inhibitor | 40 | * | 40 | * | 40 | * | 40 | * | 41 | * |
| Penicillin | Cell wall synthesis inhibitor | 21 | R | 24 | R | 24 | R | 26 | R | 22 | R |
| Ceftaroline | Cell wall synthesis inhibitor | 36 | * | 37 | S | 37 | S | 38 | S | 36 | S |
| Linezolid | Protein Synthesis Inhibitor | 38 | S | 38 | S | 37 | S | 39 | S | 38 | S |
| Doxycycline | Protein Synthesis Inhibitor | 32 | S | 34 | S | 34 | S | 36 | S | 34 | S |
| Rifampin | RNA synthesis inhibitor | 37 | S | 37 | S | 38 | S | 41 | S | 37 | S |
| TMP-SMX | DHFR inhibitor (thymidine/DNA synthesis) | 33 | S | 34 | S | 35 | S | 35 | S | 34 | S |
| Clindamycin | Protein Synthesis Inhibitor | 29 | S | 30 | S | 30 | S | 30 | S | 30 | S |
| Erythromycin | Protein Synthesis Inhibitor | 32 | S | 33 | S | 36 | S | 35 | S | 32 | S |
| Vancomycin | Cell wall synthesis inhibitor | 20 | * | 20 | * | 20 | * | 22 | * | 20 | * |
| Delafloxacin | DNA gyrase inhibitor | 51 | * | 53 | * | 52 | * | 54 | * | 52 | * |
| Chloromphenicol | Protein Synthesis Inhibitor | 25 | S | 25 | S | 25 | S | 26 | S | 25 | S |
| Synercid | Protein Synthesis Inhibitor | 30 | S | 32 | S | 32 | S | 31 | S | 32 | S |
| Ciprofloxacin | DNA gyrase inhibitor | 37 | S | 40 | S | 39 | S | 39 | S | 39 | S |
| Gentamicin | PRotein Synthesis Inhibitor | 33 | S | 35 | S | 34 | S | 35 | S | 34 | S |
| Nitrofurantoin | RNA and DNA disruption | 25 | S | 28 | S | 28 | S | 31 | S | 26 | S |
| Oxacillin | Cell wall synthesis inhibitor | 22 | S | 22 | S | 21 | S | 23 | S | 22 | S |
| Daptomycin (e-test) | Cell membrane disruptor, cell depolarizer | 0.032 | S | 0.032 | S | 0.032 | S | 0.064 | S | 0.047 | S |

| | Sp80xe | | Sp80xf | | Sp80xg | | Sp80xh | | SpMJ40.1 | | SpMJ40.2 | | Sp40xa | |
|---|---|---|---|---|---|---|---|---|---|---|---|---|---|---|
| **Antimicrobial** | **Zone (in mm)** | **S/R?** | **Zone (in mm)** | **S/R?** | **Zone (in mm)** | **S/R?** | **Zone (in mm)** | **S/R?** | **Zone (in mm)** | **S/R?** | **Zone (in mm)** | **S/R?** | **Zone (in mm)** | **S/R?** |
| Cefoxitin | 41 | * | 40 | * | 41 | * | 40 | * | 40 | * | 40 | * | 37 | * |
| Penicillin | 24 | R | 24 | R | 24 | R | 23 | R | 23 | R | 21 | R | 17 | R |
| Ceftaroline | 37 | S | 38 | S | 36 | S | 36 | S | 37 | S | 36 | S | 33 | S |
| Linezolid | 37 | S | 37 | S | 38 | S | 38 | S | 38 | S | 36 | S | 34 | S |
| Doxycycline | 34 | S | 35 | S | 34 | S | 33 | S | 38 | S | 33 | S | 31 | S |
| Rifampin | 37 | S | 37 | S | 37 | S | 38 | S | 38 | S | 38 | S | 34 | S |
| TMP-SMX | 35 | S | 34 | S | 35 | S | 34 | S | 34 | S | 34 | S | 32 | S |
| Clindamycin | 30 | S | 30 | S | 30 | S | 29 | S | 29 | S | 29 | S | 26 | S |
| Erythromycin | 33 | S | 33 | S | 32 | S | 32 | S | 33 | S | 33 | S | 29 | S |
| Vancomycin | 20 | * | 21 | * | 20 | * | 20 | * | 20 | * | 20 | * | 18 | * |
| Delafloxacin | 52 | * | 53 | * | 52 | * | 53 | * | 52 | * | 52 | * | 42 | * |
| Chloromphenicol | 25 | S | 25 | S | 25 | S | 25 | S | 26 | S | 25 | S | 24 | S |
| Synercid | 32 | S | 32 | S | 32 | S | 32 | S | 33 | S | 31 | S | 30 | S |
| Ciprofloxacin | 40 | S | 39 | S | 39 | S | 39 | S | 39 | S | 38 | S | 36 | S |
| Gentamicin | 35 | S | 34 | S | 35 | S | 33 | S | 34 | S | 32 | S | 30 | S |
| Nitrofurantoin | 27 | S | 27 | S | 27 | S | 27 | S | 26 | S | 26 | S | 24 | S |
| Oxacillin | 22 | S | 21 | S | 22 | S | 22 | S | 22 | S | 22 | S | 22 | S |
| Daptomycin (e-test) | 0.047 | S | 0.032 | S | 0.032 | S | 0.032 | S | 0.032 | S | 0.047 | S | 0.047 | S |

| | Sp40xb | | Sp40xc | | Sp40xd | | Sp40xf | | Sp40xh | | Sp40xe | | Sp40xf | |
|---|---|---|---|---|---|---|---|---|---|---|---|---|---|---|
| **Antimicrobial** | **Zone (in mm)** | **S/R?** | **Zone (in mm)** | **S/R?** | **Zone (in mm)** | **S/R?** | **Zone (in mm)** | **S/R?** | **Zone (in mm)** | **S/R?** | **Zone (in mm)** | **S/R?** | **Zone (in mm)** | **S/R?** |
| Cefoxitin | 39 | * | 36 | * | 38 | * | 38 | * | 38 | * | 40 | * | 38 | * |
| Penicillin | 22 | R | 18 | R | 21 | R | 22 | R | 21 | R | 21 | R | 20 | R |
| Ceftaroline | 36 | S | 34 | S | 34 | S | 34 | S | 34 | S | 35 | S | 35 | S |
| Linezolid | 36 | S | 35 | S | 33 | S | 35 | S | 34 | S | 36 | S | 35 | S |
| Doxycycline | 34 | S | 31 | S | 31 | S | 34 | S | 32 | S | 36 | S | 38 | S |
| Rifampin | 35 | S | 34 | S | 34 | S | 35 | S | 36 | S | 36 | S | 35 | S |
| TMP-SMX | 35 | S | 32 | S | 33 | S | 34 | S | 33 | S | 34 | S | 34 | S |
| Clindamycin | 29 | S | 26 | S | 27 | S | 28 | S | 28 | S | 28 | S | 28 | S |
| Erythromycin | 31 | S | 29 | S | 30 | S | 29 | S | 30 | S | 31 | S | 31 | S |
| Vancomycin | 20 | * | 18 | * | 19 | * | 19 | * | 20 | * | 20 | * | 20 | * |
| Delafloxacin | 50 | * | 48 | * | 49 | * | 49 | * | 49 | * | 50 | * | 50 | * |
| Chloromphenicol | 25 | S | 25 | S | 24 | S | 25 | S | 25 | S | 26 | S | 26 | S |
| Synercid | 31 | S | 30 | S | 30 | S | 30 | S | 30 | S | 31 | S | 30 | S |
| Ciprofloxacin | 38 | S | 37 | S | 36 | S | 36 | S | 36 | S | 36 | S | 37 | S |
| Gentamicin | 30 | S | 29 | S | 29 | S | 31 | S | 30 | S | 31 | S | 29 | S |
| Nitrofurantoin | 27 | S | 25 | S | 26 | S | 27 | S | 25 | S | 28 | S | 27 | S |
| Oxacillin | 23 | S | 21 | S | 22 | S | 27 | S | 22 | S | 22 | S | 22 | S |
| Daptomycin (e-test) | 0.023 | S | 0.047 | S | 0.032 | S | 0.047 | S | 0.016 | S | 0.023 | S | 0.023 | S |

| | WT *S. schleiferi* | | Ss40xa | | Ss40xd | | Ss40xg | | Ss40xe | |
|---|---|---|---|---|---|---|---|---|---|---|
| **Antimicrobial** | **Zone (in mm)** | **S/R?** | **Zone (in mm)** | **S/R?** | **Zone (in mm)** | **S/R?** | **Zone (in mm)** | **S/R?** | **Zone (in mm)** | **S/R?** |
| Cefoxitin | 36 | * | 39 | * | 35 | * | 36 | * | 36 | * |
| Penicillin | 50 | S | 48 | S | 58 | S | 46 | S | 48 | S |
| Ceftaroline | 42 | * | 42 | * | 42 | * | 40 | * | 40 | * |
| Linezolid | 33 | S | 32 | S | 32 | S | 36 | S | 32 | S |
| Doxycycline | 33 | S | 32 | S | 31 | S | 38 | S | 31 | S |
| Rifampin | 37 | S | 36 | S | 36 | S | 38 | S | 35 | S |
| TMP-SMX | 29 | S | 27 | S | 27 | S | 28 | S | 28 | S |
| Clindamycin | 29 | S | 29 | S | 27 | S | 29 | S | 26 | S |
| Erythromycin | 30 | S | 28 | S | 29 | S | 30 | S | 27 | S |
| Vancomycin | 19 | * | 19 | * | 19 | * | 20 | * | 18 | * |
| Delafloxacin | 42 | * | 40 | * | 41 | * | 41 | * | 40 | * |
| Chloromphenicol | 25 | S | 26 | S | 25 | S | 24 | S | 23 | S |
| Synercid | 30 | S | 30 | S | 31 | S | 29 | S | 28 | S |
| Ciprofloxacin | 32 | S | 31 | S | 31 | S | 30 | S | 30 | S |
| Gentamicin | 33 | S | 30 | S | 29 | S | 30 | S | 28 | S |
| Nitrofurantoin | 25 | S | 25 | S | 24 | S | 24 | S | 23 | S |
| Oxacillin | 25 | S | 26 | S | 25 | S | 26 | S | 25 | S |
| Daptomycin (e-test) | 0.047 | S | 0.047 | S | 0.032 | S | 0.032 | S | 0.047 | S |

| | Ss80xa | | Ss80xe | | Ss80xf | | Ss80xg | |
|---|---|---|---|---|---|---|---|---|
| **Antimicrobial** | **Zone (in mm)** | **S/R?** | **Zone (in mm)** | **S/R?** | **Zone (in mm)** | **S/R?** | **Zone (in mm)** | **S/R?** |
| Cefoxitin | 36 | * | 34 | * | 34 | * | 35 | * |
| Penicillin | 50 | S | 48 | S | 48 | S | 48 | S |
| Ceftaroline | 42 | * | 42 | * | 42 | * | 42 | * |
| Linezolid | 35 | S | 34 | S | 35 | S | 40 | S |
| Doxycycline | 33 | S | 31 | S | 33 | S | 35 | S |
| Rifampin | 37 | S | 35 | S | 38 | S | 37 | S |
| TMP-SMX | 27 | S | 29 | S | 29 | S | 30 | S |
| Clindamycin | 29 | S | 28 | S | 31 | S | 31 | S |
| Erythromycin | 32 | S | 30 | S | 35 | S | 35 | S |
| Vancomycin | 19 | * | 18 | * | 21 | * | 20 | * |
| Delafloxacin | 42 | * | 41 | * | 43 | * | 43 | * |
| Chloromphenicol | 25 | S | 24 | S | 28 | S | 27 | S |
| Synercid | 30 | S | 30 | S | 32 | S | 33 | S |
| Ciprofloxacin | 32 | S | 31 | S | 32 | S | 34 | S |
| Gentamicin | 30 | S | 29 | S | 31 | S | 33 | S |
| Nitrofurantoin | 24 | S | 24 | S | 26 | S | 27 | S |
| Oxacillin | 25 | S | 26 | S | 26 | S | 27 | S |
| Daptomycin (e-test) | 0.047 | S | 0.032 | S | 0.047 | S | 0.047 | S |

Table S2. Minimum inhibitory concentrations (MIC) for values for selected antistaphylococcals against POM-ERJ resistant staphylococci, R1-R3. Displayed are the mean ± SD of three independent biological experiments performed in technical duplicate. In some cases, SD is listed as N/A as MIC values are discrete measurements and each replicate provided the same measurement, hence there is no variability.

| Compound | Wild-Type | | R1 | | R2 | | R3 | |
|---|---|---|---|---|---|---|---|---|
| | MIC (μM) | SD | MIC (μM) | SD | MIC (μM) | SD | MIC (μM) | SD |
| FSM | 6.3 | N/A | 5.2 | 1.6 | 5.2 | 1.6 | 5.2 | 1.6 |
| FR-900098 | 500 | N/A | 500 | N/A | 500 | N/A | 500 | N/A |
| BOM-ERJ | 7.9 | 4.6 | 27 | N/A | 27 | N/A | 27 | N/A |
| POM-ERJ | 0.50 | N/A | 180 | N/A | 37 | 13 | 180 | N/A |
| Cefditoren Sodium | 0.74 | 0.54 | 0.55 | 0.27 | 0.59 | 0.23 | 0.59 | 0.23 |
| Cefditoren Pivoxil | 0.76 | N/A | 12 | N/A | 6.0 | N/A | 10 | 3.1 |
| HEX | | | | | | | | |
| POM-HEX | 3.1 | N/A | 100 | N/A | 50 | N/A | 100 | N/A |
| Mupirocin | 0.010 | 0.0041 | 0.010 | 0.0041 | 0.010 | 0.0041 | 0.0092 | 0.0032 |

Table S3. Single Nucleotide Polymorphisms identified via whole-genome sequencing.

| Strain | Species | Base | WT Allele | SNP | Read Depth | Gene | Annotation |
|---|---|---|---|---|---|---|---|
| sp40xa | *S. pseudintermedius* | 45418 | G | A | 5 | SPSE_0038 | P-type Copper Transporter |
| sp40xa | *S. pseudintermedius* | 1282386 | A | T | 219 | SPSE_1252 | Hydroxyacylglutathione hydrolase |
| sp40xa | *S. pseudintermedius* | 2227530 | C | A | 562 | SPSE_2164 | Putative glutamyl-endopeptidase |
| sp40xa | *S. pseudintermedius* | 2228995 | C | A | 519 | SPSE_2165 | Putative glutamyl-endopeptidase |
| sp40xb | *S. pseudintermedius* | 45418 | G | A | 5 | SPSE_0038 | P-type Copper Transporter |
| sp40xb | *S. pseudintermedius* | 1282347 | C | A | 230 | SPSE_1252 | Hydroxyacylglutathione hydrolase |
| sp40xc | *S. pseudintermedius* | 45418 | G | A | 5 | SPSE_0038 | P-type Copper Transporter |
| sp40xc | *S. pseudintermedius* | 1110010 | A | T | 75 | SPSE_1082 | Transposase |
| sp40xc | *S. pseudintermedius* | 2227530 | C | A | 722 | SPSE_2164 | Putative glutamyl-endopeptidase |
| sp40xd | *S. pseudintermedius* | 1282655 | C | T | 358 | SPSE_1252 | Hydroxyacylglutathione hydrolase |
| sp40xe | *S. pseudintermedius* | 1282745 | G | T | 406 | SPSE_1252 | Hydroxyacylglutathione hydrolase |
| sp40xe | *S. pseudintermedius* | 1651160 | C | A | 493 | SPSE_1610 | Putative oxidoreductase |
| sp40xf | *S. pseudintermedius* | 1282565 | C | T | 283 | SPSE_1252 | Hydroxyacylglutathione hydrolase |
| sp40xf | *S. pseudintermedius* | 2228995 | C | A | 618 | SPSE_2165 | Putative glutamyl-endopeptidase |
| sp40xg | *S. pseudintermedius* | 2228995 | C | A | 526 | SPSE_2165 | Putative glutamyl-endopeptidase |
| sp40xh | *S. pseudintermedius* | 1282347 | C | A | 279 | SPSE_1252 | Hydroxyacylglutathione hydrolase |
| sp40xh | *S. pseudintermedius* | 1651160 | C | A | 492 | SPSE_1610 | Putative oxidoreductase |
| sp40xh | *S. pseudintermedius* | 2227530 | C | A | 611 | SPSE_2164 | Putative glutamyl-endopeptidase |
| sp80xa | *S. pseudintermedius* | 1282655 | C | T | 328 | SPSE_1252 | Hydroxyacylglutathione hydrolase |
| sp80xb | *S. pseudintermedius* | 2227530 | C | A | 679 | SPSE_2164 | Putative glutamyl-endopeptidase |
| sp80xc | *S. pseudintermedius* | 1282655 | C | T | 372 | SPSE_1252 | Hydroxyacylglutathione hydrolase |

| Strain | Species | Base | WT Allele | SNP | Read Depth | Gene | Annotation |
|---|---|---|---|---|---|---|---|
| sp80xd | *S. pseudintermedius* | 2228995 | C | A | 586 | SPSE_2165 | Putative glutamyl-endopeptidase |
| sp80xe | *S. pseudintermedius* | 668455 | T | G | 5 | SPSE_0610 | RNA-directed DNA polymerase |
| sp80xe | *S. pseudintermedius* | 1282874 | G | A | 292 | SPSE_1252 | Hydroxyacylglutathione hydrolase |
| sp80xe | *S. pseudintermedius* | 1651160 | C | A | 415 | SPSE_1610 | Putative oxidoreductase |
| sp80xf | *S. pseudintermedius* | 668455 | T | G | 6 | SPSE_0610 | RNA-directed DNA polymerase |
| sp80xf | *S. pseudintermedius* | 1282386 | A | T | 327 | SPSE_1252 | Hydroxyacylglutathione hydrolase |
| sp80xf | *S. pseudintermedius* | 2227530 | C | A | 545 | SPSE_2164 | Putative glutamyl-endopeptidase |
| sp80xg | *S. pseudintermedius* | 1282347 | C | A | 280 | SPSE_1252 | Hydroxyacylglutathione hydrolase |
| sp80xg | *S. pseudintermedius* | 2227530 | C | A | 613 | SPSE_2164 | Putative glutamyl-endopeptidase |
| sp80xh | *S. pseudintermedius* | 1282385 | G | T | 35 | SPSE_1252 | Hydroxyacylglutathione hydrolase |
| sp80xh | *S. pseudintermedius* | 2228043 | A | C | 174 | SPSE_2164 | Putative glutamyl-endopeptidase |
| sp80xh | *S. pseudintermedius* | 2227746 | C | A | 226 | SPSE_2164 | Putative glutamyl-endopeptidase |
| sp80xh | *S. pseudintermedius* | 2227738 | G | A | 242 | SPSE_2164 | Putative glutamyl-endopeptidase |
| sp80xh | *S. pseudintermedius* | 2227731 | C | A | 251 | SPSE_2164 | Putative glutamyl-endopeptidase |
| sp80xh | *S. pseudintermedius* | 2227735 | G | C | 255 | SPSE_2164 | Putative glutamyl-endopeptidase |
| sp80xh | *S. pseudintermedius* | 2227530 | C | A | 579 | SPSE_2164 | Putative glutamyl-endopeptidase |
| sp80xh | *S. pseudintermedius* | 2228995 | C | A | 589 | SPSE_2165 | Putative glutamyl-endopeptidase |
| spmj401 | *S. pseudintermedius* | 1282794 | G | T | 225 | SPSE_1252 | Hydroxyacylglutathione hydrolase |
| spmj401 | *S. pseudintermedius* | 2228995 | C | A | 520 | SPSE_2165 | Putative glutamyl-endopeptidase |
| spmj402 | *S. pseudintermedius* | 1282329 | G | T | 240 | SPSE_1252 | Hydroxyacylglutathione hydrolase |
| spmj402 | *S. pseudintermedius* | 2227530 | C | A | 597 | SPSE_2164 | Putative glutamyl-endopeptidase |
| ss40xa | *S. schleiferi* | 925052 | G | A | 526 | gatA | Galactitol PTS system EIIA component |

| Strain | Species | Base | WT Allele | SNP | Read Depth | Gene | Annotation |
|---|---|---|---|---|---|---|---|
| ss40xd | *S. schleiferi* | 925052 | G | A | 465 | gatA | Galactitol PTS system EIIA component |
| ss40xe | *S. schleiferi* | 178904 | G | T | 543 | LH95_00765 | Putative Tributyrin Esterase |
| ss40xf | *S. schleiferi* | 124153 | G | T | 564 | murQ | N-acetylmuramic acid 6-phosphate etherase |
| ss40xg | *S. schleiferi* | 971929 | G | A | 18 | LH95_04455 | tRNA Val |
| ss40xg | *S. schleiferi* | 971915 | A | T | 43 | LH95_04455 | tRNA Val |
| ss40xg | *S. schleiferi* | 971919 | G | T | 43 | LH95_04455 | tRNA Val |
| ss40xg | *S. schleiferi* | 971911 | C | A | 45 | LH95_04455 | tRNA Val |
| ss40xg | *S. schleiferi* | 971900 | G | T | 155 | LH95_04455 | tRNA Val |
| ss40xg | *S. schleiferi* | 2095855 | T | C | 541 | LH95_09910 | Potassium-transporting ATPase potassium-binding subunit |
| ss80xa | *S. schleiferi* | 1282737 | C | A | 347 | LH95_06060 | Hydroxyacylglutathione hydrolase |
| ss80xe | *S. schleiferi* | 1282513 | G | A | 424 | LH95_06060 | Hydroxyacylglutathione hydrolase |
| ss80xe | *S. schleiferi* | 1291886 | G | A | 386 | LH95_06115 | Lipoyl(octanoyl) transferase |
| ss80xf | *S. schleiferi* | 1282335 | C | T | 289 | LH95_06060 | Hydroxyacylglutathione hydrolase |
| ss80xg | *S. schleiferi* | 1291886 | G | A | 294 | LH95_06115 | Lipoyl(octanoyl) transferase |
| ss80xh | *S. schleiferi* | 1319671 | G | T | 406 | LH95_06255 | Ribonuclease Z |
| ss80xh | *S. schleiferi* | 1478053 | C | G | 458 | LH95_07010 | Unknown function |

Table S4. Primers used during this study.

| **Primer Sequence** | **Gene** | **Species** | **Purpose** |
|---|---|---|---|
| ATATCGCTGCATTAGATGATG | SPSE_1252(gloB) | S. pseudintermedius | Sequencing |
| AGGCACATCATATCGTGTTAG | SPSE_1252(gloB) | S. pseudintermedius | Sequencing |
| TGCTGCATTCCTTCATCAAGTG | SPSE_1252(gloB) | S. pseudintermedius | Sequencing |
| CCGTTCAATAAAGGGCTCGATC | SPSE_1252(gloB) | S. pseudintermedius | Sequencing |
| TGATGAATTTGCAGTAGTGGGC | SPSE_1252(gloB) | S. pseudintermedius | Sequencing |
| TTAACCTTGACCGTCTAAAAACG | SPSE_1252(gloB) | S. pseudintermedius | Sequencing |
| CTGGGCAAAAGCAGTATTGACAGGC | SPSE_2164 | S. pseudintermedius | Sequencing |
| GTTAGCGGTTTCACAGATGCC | SPSE_2164 | S. pseudintermedius | Sequencing |
| AGTTCGGGTTATTCATCCTAACACC | SPSE_2164 | S. pseudintermedius | Sequencing |
| CGTCAACTGTCGCATTAACTGC | SPSE_2164 | S. pseudintermedius | Sequencing |
| TAATTTGCGTTTTTGTTAACCC | SPSE_2164 | S. pseudintermedius | Sequencing |
| GTTAGCGGTTTCACAGATGCC | SPSE_2164 | S. pseudintermedius | Sequencing |
| ATGAAAATTTCCTACCTGACTTTAG | LH95_06060(gloB) | S. schleiferi | Sequencing |
| ATTTTTTTTGATTCTGAAGATGGG | LH95_06060(gloB) | S. schleiferi | Sequencing |
| ATGCACAGCCTACTGCGATCGAAG | LH95_06060(gloB) | S. schleiferi | Sequencing |
| TTACTCATGCACATTTTGATC | LH95_06060(gloB) | S. schleiferi | Sequencing |
| AATGCCCAGGTGTATGCAATGC | LH95_06060(gloB) | S. schleiferi | Sequencing |
| TTAGCCATGAAGATAAGGATTC | LH95_06060(gloB) | S. schleiferi | Sequencing |
| GGAGTGAGTATTTTGGCACG | LH95_00765 | S. schleiferi | Sequencing |
| AGGATCAAAAGGTGTCCCCACAAC | LH95_00765 | S. schleiferi | Sequencing |
| GCGGTGATTATGCCCAATGCAGACC | LH95_00765 | S. schleiferi | Sequencing |
| TTATACCATCTCACGCGTATGATGG | LH95_00765 | S. schleiferi | Sequencing |
| CTCACCACCACCACCACCATATGAAAATTTCCTACCTGACTTTAG | LH95_06060(gloB) | S. schleiferi | LIC Cloning |
| ATCCTATCTTACTCACTTAGCCATGAAGATAAGGATTC | LH95_06060(gloB) | S. schleiferi | LIC Cloning |
| CTCACCACCACCACCACCATATGAATATTTCTAATCTTACTTTAG | SPSE_1252(gloB) | S. pseudintermedius | LIC Cloning |
| ATCCTATCTTACTCACTTAACCTTGACCGTCTAAAAACG | SPSE_1252(gloB) | S. pseudintermedius | LIC Cloning |
